# Cloning human dental pulp cells and studying inter-clone diversity

**DOI:** 10.1101/703280

**Authors:** Linna Guo, Ziang Zou, Ming Yan, Marcus Freytag, Reinhard E Friedrich, Lan Kluwe

## Abstract

Heterogeneity within a putative stem cell population presents a challenge for studies and applications of such cells. Cloning may provide a strategy for reducing heterogeneity. However, previous studies have the weakness in reliability of single-cell-origin of the colonies. The present study aims to apply an alternative method to obtain clonal dental pulp cells with increased reliability of single cell origin. Dental pulp cells were cultured from 13 human wisdom teeth. Primary cultures of 3 human benign tumors were included as comparison. Cells were seeded into wells of a 96-plate at a mean density of 1 cell/well. On the next day, wells were inspected one by one to identify wells with single cells which were followed for 3 weeks. Survived clones were expanded and further characterized. Single cells were observed in all cases, the number of single-cell-wells varied from 16 to 33. Three weeks later, survived and grown clonal cells were observed in 10 to 29 wells, giving surviving rates of 33-91%. By contrast, though single tumor cells were also observed, none of them survived. Expanded clones exhibited diversity in viability and osteogenic differentiation which also differed from their parental cells. Seeding cells at clonal density into physically separated compartments like wells in a 96 plate and comprehensive observation provides a practical strategy for increasing reliability of single-cell origin of clonal cells. Cellular heterogeneity seems to be an intrinsic feature of dental pulp cells.

## Introduction

Human dental pulp cells exhibit stem/progenitor cell like features such as clonal growth, high proliferation rate and capacity of differentiating into multiple cell types including adipocytes, osteocytes and chondrocytes [1, 2]. Because of their easy and non-invasive accessibility [3, 4], dental pulp cells provide a valuable source of stem/progenitor cells for tissue engineering and regenerative medicine.

However, as majority of primary cells, dental pulp cells are heterogeneous among donors and also within cells from a single tooth [5, 6]. This kind of cellular heterogeneity poses an obstacle in expectation and in reproducibility of experiments and in further development of their application. Therefore, strategies have be conceived and practiced in order to obtain more homogeneous cells and to enrich stem/progenitor cells. For example, sorting cells based on their surface markers using flow cytometry is one well established method. However, specificities of surface markers for stem/progenitor cells are still controversial which also likely vary depending on cells passage, density, cultural conditions and donor teeth. Moreover, the issue is yet to be comprehensively addressed if and to which extent purified or enriched subpopulation may become heterogeneous as they are expanded.

Clonal growth is a major feature of stem/progenitor cells. A hierarchical structure is hypothesized for mesenchymal stem cells, that is, mother stem cells possess highest self-renewal ability and greatest potential to form colonies, and that single cells which can form colonies are likely the mother stem cells [5]. Indeed, low density seeding has been applied to stem/progenitor cell populations including periodontal ligament cells, bone marrow mesenchymal stem cells and cancer stem in order to select mother stem cells [7–9]. However, as argued in a recent review [10], a colony may contain two or more cells in proximity and colonies may fuse together. As emphasized in this review, for addressing the issue of intra-clonal heterogeneity, it is critical first to confirm thoroughly that the cells in question indeed originate from single cells.

Our previous study demonstrated the survival of dental pulp cells when seeded at clonal density in wells of 96 plate [11]. However, individual clones were not thoroughly checked and followed from the first day. In the present study, we seeded dental pulp cells into wells of a 96 plate at a mean density of 1 cell/well and followed single cells from day 1 to day 21. Physical separation of the wells prevents cells moving together and fusion of colonies. Large number of compartment (96 wells) enables statistical estimation assuming a Poisson distribution which is that is 35 (37%) wells from a 96-plate will contain no cells, 35 (37%) will contain single cells and only 16 will contain 2 or more in each (Fig 1). As a comparison, primary cultures of 3 human benign tumors (plexiform neurofibromas) were also included. Survived clonal dental pulp cells were expanded and further characterized regarding their growth and differentiation properties.

**Fig 1.**
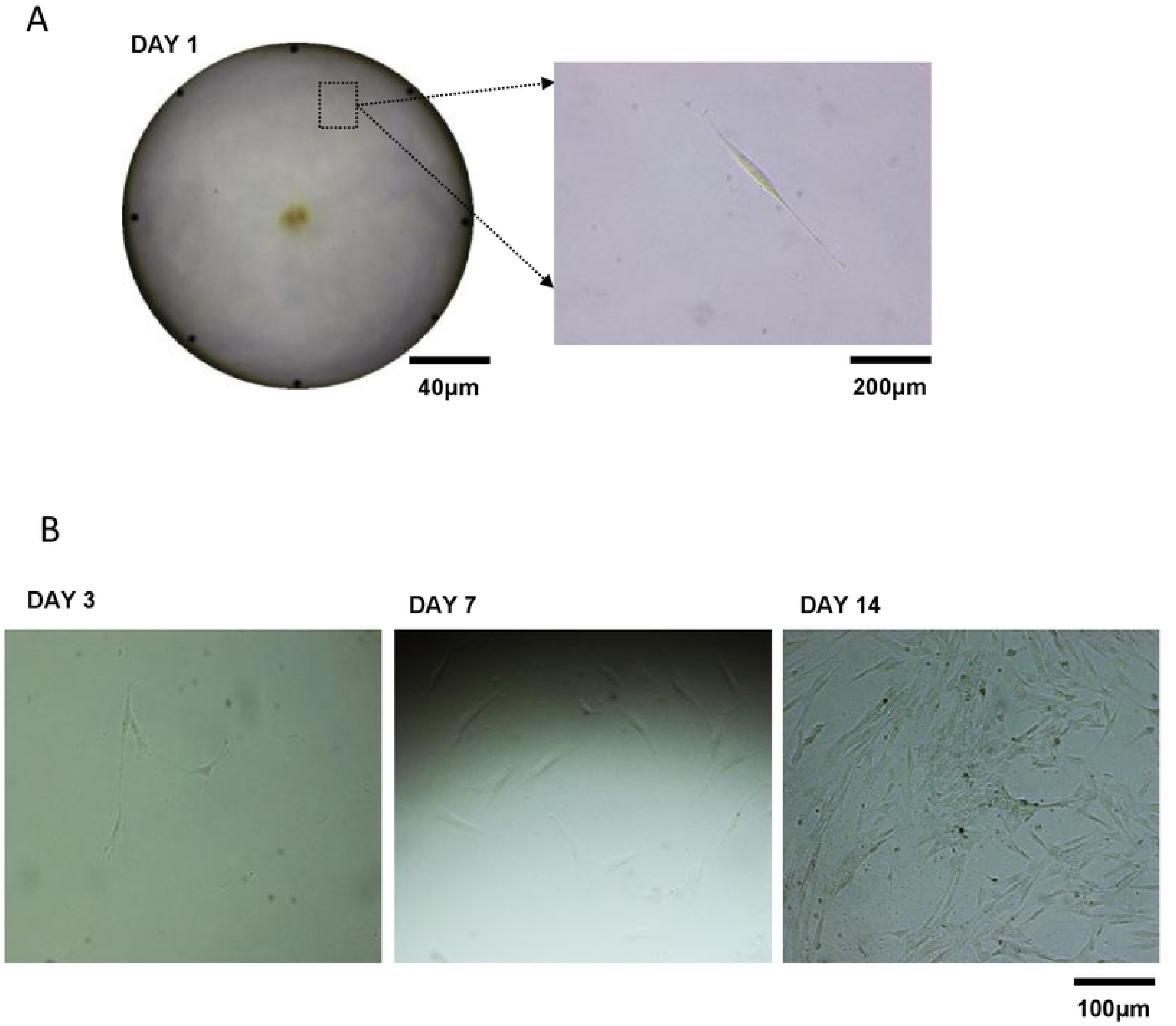
An example of a single cell at seeding and its subsequent clonal growth. Dental pulp cells derived from wisdom tooth were seeded into wells of a 96-plate at density of 1 cell/well. On the next day (day 1), wells were checked one by one. **A:** a representative well with a single cell. **B:** clonal growth of this single cell on day 3, 7 and 14.

## Materials and methods

### Primary cell culture

Human dental pulp cells were cultured from a total of 13 anonymized wisdom teeth of adults (18–51 years old) which were extracted at our Department of Oral and Maxillofacial Surgery, Eppendorf Medical Center Hamburg-Eppendorf, Germany. The outgrowth method was used to culture dental pulp cells [11]. Briefly, the teeth were broken with a hammer, the pulp tissues were taken out and cut into small pieces which were placed onto cultural surface in wells of 6-well plates and cultured in MEM with 15% fetal bovine serum and antibiotics. Some of the dental pulp cells were frozen stored at −80°C for short periods.

As comparison, primary cells were also cultured from 3 anonymized plexiform neurofibromas which are benign human tumors of the peripheral nerve [12].

The specimens were recovered from bio-waste and completely anonymized according the local privacy protection regulation. The study protocol was reported to the corresponding local authority. All patients signed written informed consent forms for using of the specimen.

### Low-density seeding

Primary dental pulp cells, frozen stored dental pulp cells and the comparative human primary tumor cells were harvested upon sub-confluence, and re-suspended at the density of 1 cell/100μl. To each well of a 96-well plate, 100μl of this suspension was added, giving an average of 1 cell/well. The distribution of the cells in the wells should follow the Poisson principle [11] which predicts 35 wells of a 96-plate having no cells, 35 wells having single cells, 18 wells having 2 cells and the rest 8 wells having more than 2 cells. Cells from each donor were seeded into 96 wells of one plate.

On the next day, the wells of a plate were checked one by one under a microscope and the number of attached cells in each well were recorded. Wells containing single cell were further followed over a period of 3 weeks. At the end of the follow-up period, number of wells containing survived and grown cells were recorded. For each of the 13 teeth, 3 randomly selected clonal cells (in 3 different wells) were further expanded in 6-well plates.

Viability of expanded clonal pulp cells and the parental pulp cells was measured using a CellTiter 96®Aqueous Non-Radioactive Cell Proliferation Assay Kit (MTS assay, Promega, USA) and differentiation of the cells were carried out as following.

### Multipotent differentiation

Cells were expanded seeded in wells of a 24-well plate and grew to approximately 80% confluence. Differentiation was then initiated by changing the media to DMEM/Hams F12 with 10% human serum supplemented with components for adipo-, osteo- and chondral-induced medium respectively:

- Adipo-differentiation: 1µM dexamethasone, 0.5mM IBMX, 0.2mM indomethacin and 10 µM insulin
- Osteogenic differentiation: 50 µM ascorbic acid 2-phosphate and 10 mM β-Glycerophosphate
- Chondrogenic differentiation: 10 µM insulin, 50 nM ascorbic acid 2-phosphate and 6.25 µg/ml Transferrin and 10ng/mL TGF-β

The media were refreshed every other day and the differentiation was continued for three weeks. At the end of the differentiation, the cells were stained with Oil-Red for oil droplets, Alizarin Red for mineral sedimentation and Alcian Blue for cartilage formation, respectively. Osteogenic differentiation was additionally quantified by measuring alkaline phosphatase activity using a p-Nitrophenyl phosphate assay [13]. The enzyme activity values were normalized against viability of the cells.

## Results

### Single cells after seeding

From all 13 wisdom teeth, vital dental pulp cells were successfully obtained. Cells derived from each tooth were seeded into wells of one 96-well plate at an average density of 1 cell/well. On the next day, wells containing single cells were found in all cases (an example in Fig 1A). These single cells attached to the surface and appeared vital. In the example of cells derived from one wisdom tooth in Fig 2, 58 wells contained no cell, 28 contained single cells and the rest 10 wells contained 2 or more cells. Across 13 plates for dental pulp cells from 13 wisdom teeth, numbers of wells containing single cells varied from 16 to 33 (Fig 3A).

**Fig 2.**
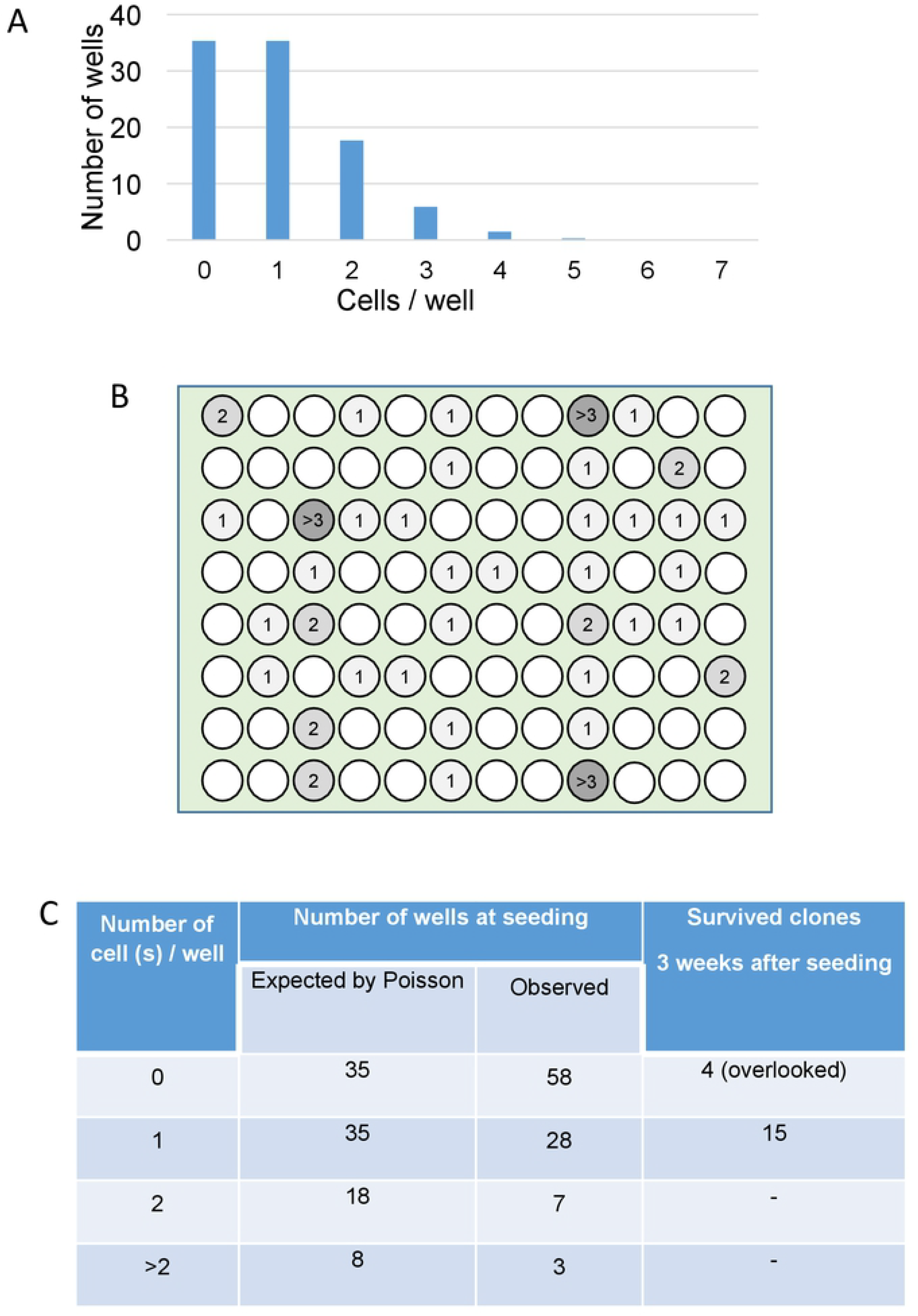
Distribution of the cells in the wells of a 96-plate. **A**: Poisson distribution showing expected number of wells containing 0, 1, 2 and >2 cells. **B**: an illustration of an example of observed distribution of cells in a 96-plate. Wells contained no cell were marked in white, wells contained 1, 2 and >2 cells were labeled with the number and illustrated in light, median and heavy grey. **C**: summary of the distribution of cells at seeding in this plate. Survived clones and overlooked clones were also given in the last column. The “4 (overlooked)” wells were initially recorded as having no cells which however turned out to have survived and grown cells later, indicating that these 4 wells had cells at seeding which were overlooked, giving a frequency of false negative of 8%.

**Fig 3.**
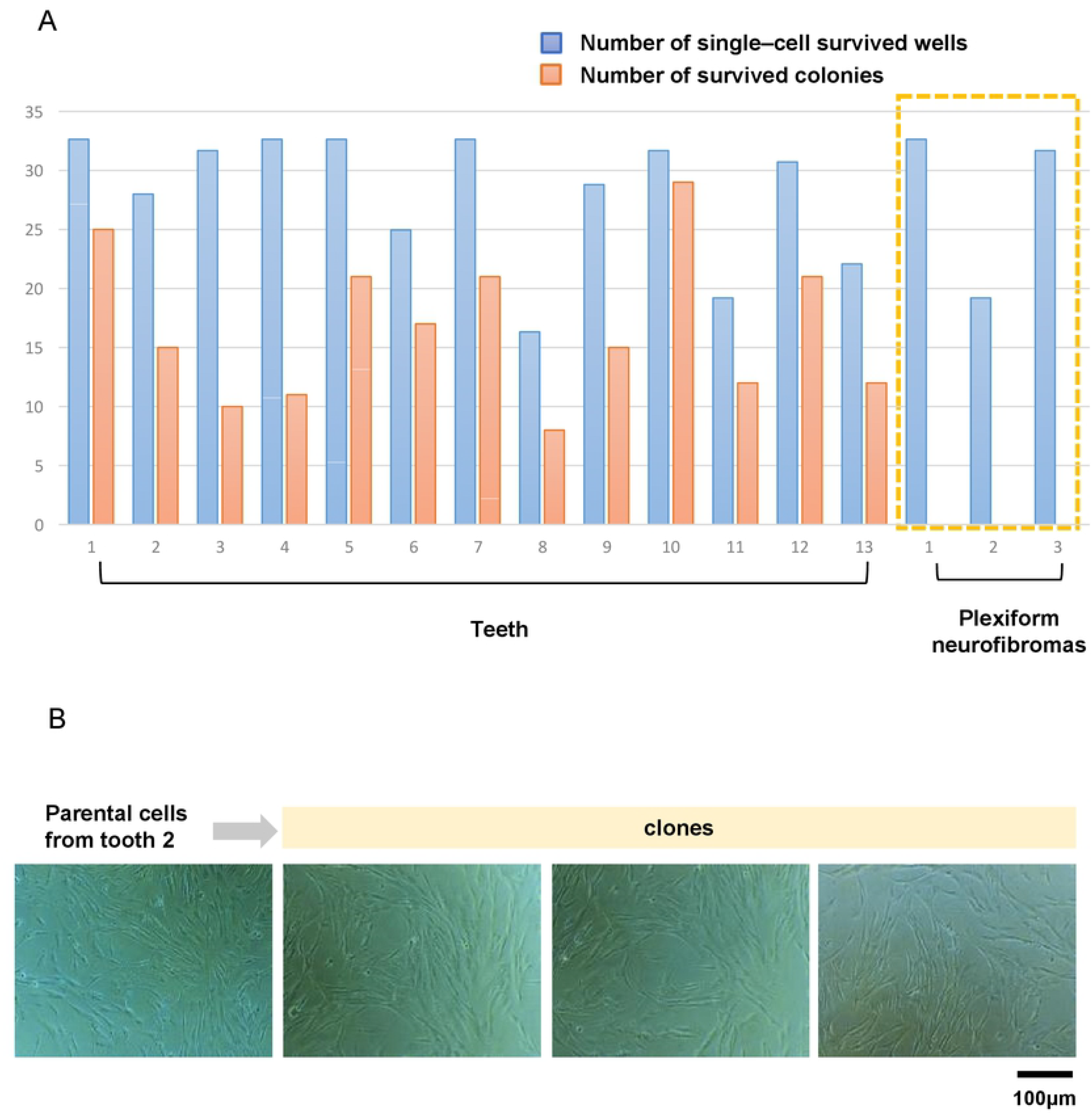
Frequency of single cells at seeding and survived single cells after 3 weeks. **A:** Blue bars represent number of wells containing single cells at seeding; orange bars represent number of survived single cells. Each pair of blue/orange bars is for cells from one donor tooth. The counting was carried out for cells of a total of 13 donor teeth and for tumor cells from 3 human benign tumors. Note that though single tumor cells were also observed at seeding, all of them died or terminated growth after a few days.

Also, for the 3 tumors, single tumor cells were found one day after the seeding at compatible frequency as in the case of the dental pulp cells. Also these single tumor cells attached to the surface and appeared vital.

The same experiment was further carried out for dental pulp cells which have been once frozen stored at −80°C. Vital single cells were observed at compatible rate as those without being frozen and thawed.

Wells with single cells were monitored every two days and followed up for 3 weeks. Some of the single dental pulp cells started growing readily on day 3 (Fig 1B). However, some other single cells died during the follow-up period. All tumor cells died few days later.

### Survival and clonal growth of single cells

Three weeks later, some single cells grew more than 30 cells which were counted as survived clones and were found for all the dental pulp cells from the 13 donor teeth. Number of clones varied from 10 to 29 and the surviving rates (number of survived clones divided by number of single cells at seeding) varied from 33% to 91% (Fig 3A). Survived clones were also obtained for dental pulp cells which have been frozen and thawed, though the surviving rate is somehow lower (25-75%).

By contrast, all tumor cells seeded at clonal density gradually died several days after the seeding (Fig 3A). Consequently, no clones were obtained from any of the 3 primary tumors.

To assess the possibility of overlooking cells, we selected 3 cases of pulp cells and rechecked the 51, 58 (Fig 2C) and 49 wells in which were recorded as having no cell at seeding, respectively. Indeed, cells were found in 3, 4 and 5 wells in each case, giving the frequency of false negative of 6% (3 in 51), 7% (4 in 58, Fig 2C) and 8% (5 in 49), respectively.

### Diversity among clones

For each of the 13 teeth, 3 clones were randomly selected and expanded for further characterization. No apparent difference in morphology was observed for dental pulp cells in different clones and their parental cells (Fig 3B).

For cells derived from each donor tooth, viability varied among cells of different clones and the clonal cells were differed from their parental mixed pulp cells. In some cases, the parental cells exhibited high viability than the 3 randomly selected clones. However, in other cases, clones exhibited higher viability than the parental cells. No straightforward rule can be speculated. Examples were given for cells from 3 teeth (Fig 4).

**Fig 4.**
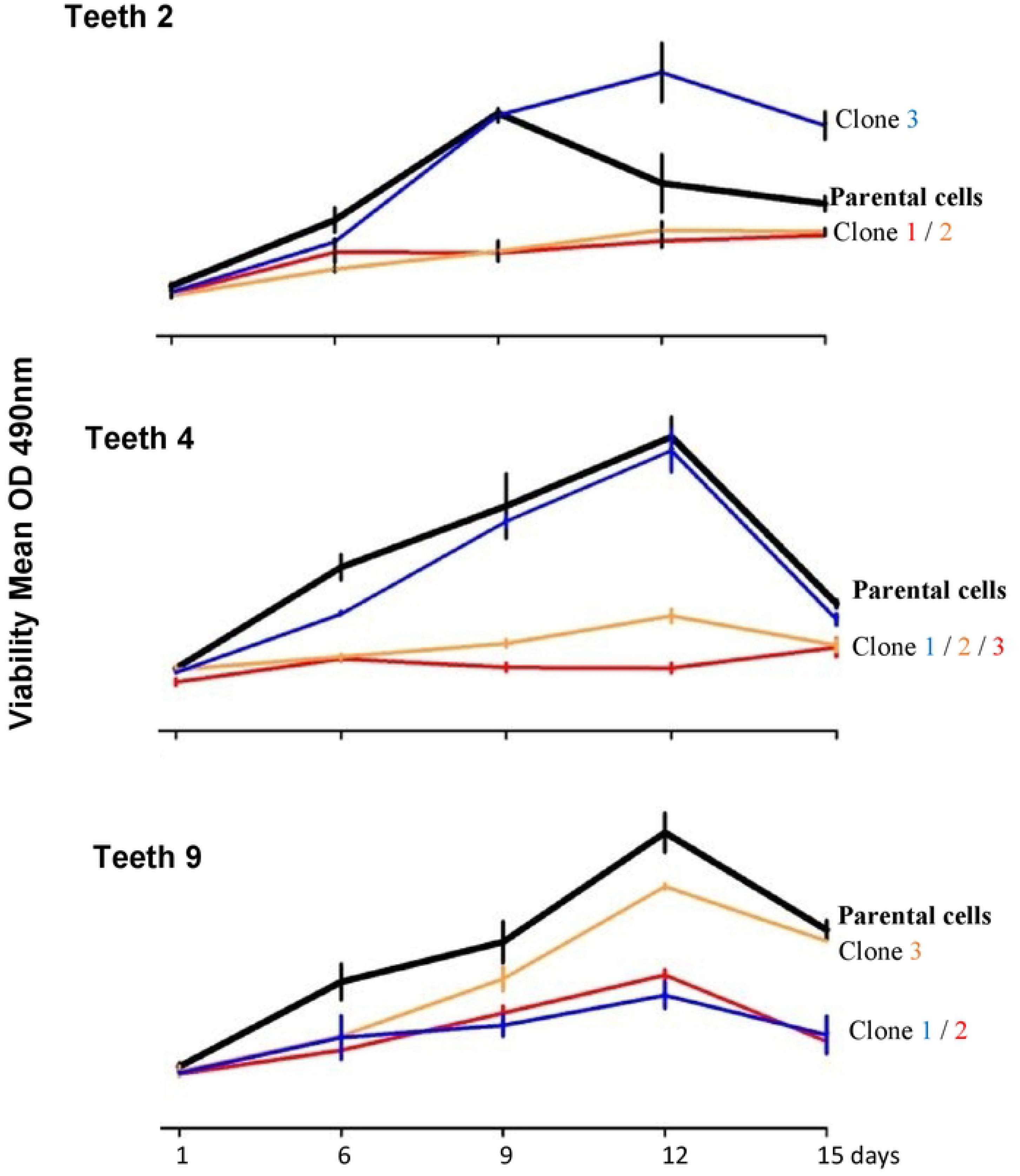
Viability of expanded clonal cells and their parental dental pulp cells. Examples were given for cells of 3 donor teeth. Viability varied among clones and between clones and their parental cells.

Adipogenic, osteogenic, and chondrogenic differentiation were successful in all expanded clonal cells and in the parental pulp cells from all the 13 donor teeth (an example in Fig 5A, C, E). These differentiations were not observed in human gingival fibroblasts under the same induction condition (Fig 5B, D, F), suggesting that the differentiation capacity is a specific feature of the dental pulp cells. Also within clones, differentiation was not homogeneous, suggesting certain kind of intra-clonal diversity. No apparent difference was observed among clones and neither between clones and their parental cells.

**Fig 5.**
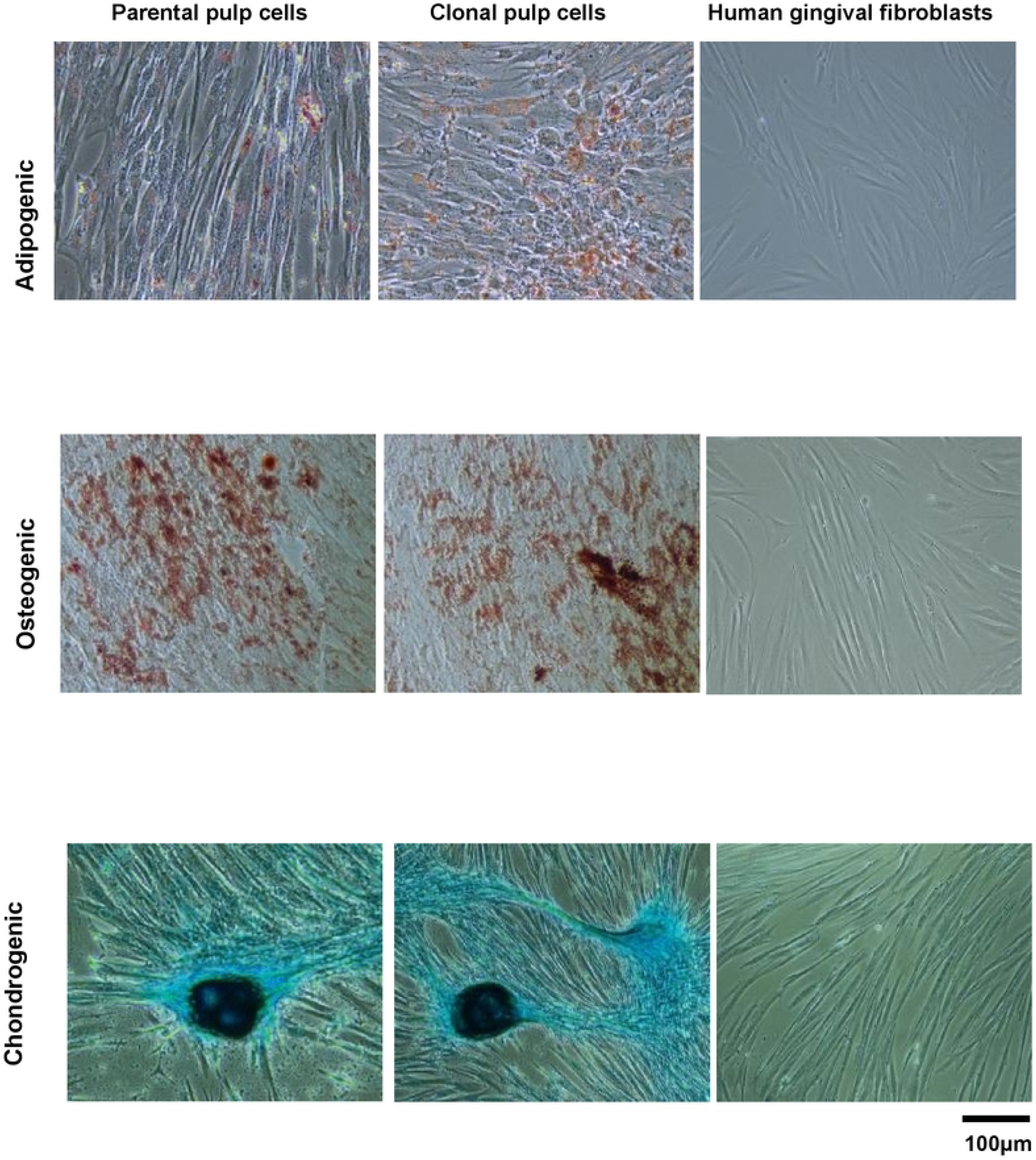
An example of differentiation of parental and clonal dental pulp cells from the donor tooth No.2. Human gingival fibroblasts were subjected to the same differentiation-induction as a negative control. Adipose differentiation was visualized by staining lipid droplets with Oil Red O. Osteogenic differentiation was visualized by staining mineral sediments with Alizarin Red. Chondrogenic differentiation was visualized by staining the chondrogenic matrix with Alcian Blue.

Osteogenic differentiation was further quantified by measuring alkaline phosphatase activity of the cells which varied among clones and between clones and their parental cells (Fig 6). Some clones had higher activity than their parental cells but there were also clones which had lower activity, also within clones derived from an identical donor tooth.

**Fig 6.**
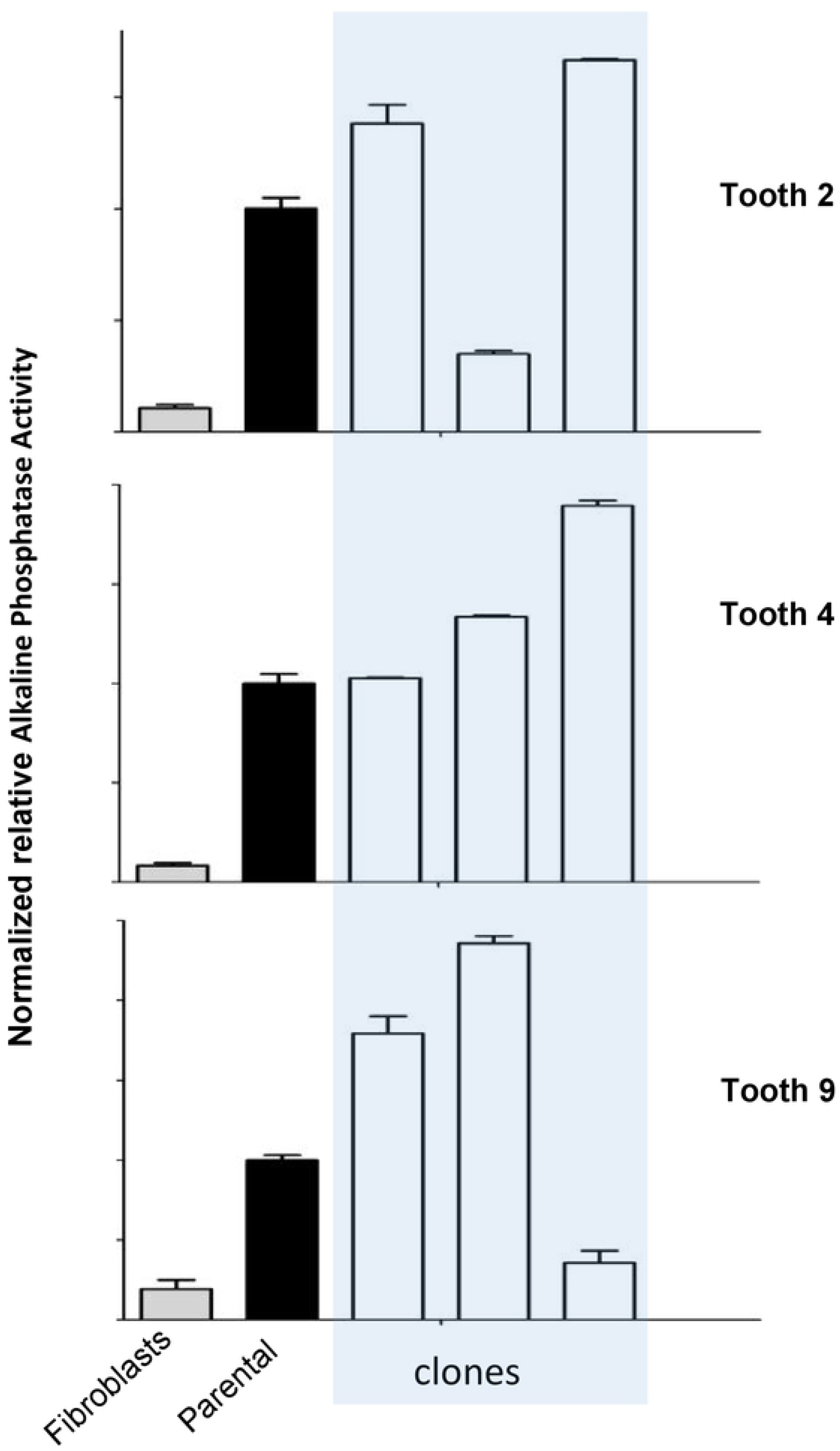
Osteogenic differentiation was quantified by measuring alkaline phosphatase activity in clonal and parental dental pulp cells from 3 donor teeth. Only background level of enzyme activity was measured in human gingival fibroblasts which were included as a negative control (grey bars). Black bars represent alkaline phosphatase activities in parental pulp cells. White bars represent the enzyme activities in different clones.

## Discussion

The present study was designed to obtain clonal cells from primary human dental pulp cells with special effort on increasing the reliability of single-cell origin of the clones. Several lines of evidence and considerations suggest that majority of our clones originate from single cells:

1. Seeding cells at clonal density into wells of a 96 plate provides a physical separation which prevents cells in proximity moving together and prevent fusion of colonies.
2. Poisson distribution predict approximately 1/3 of the wells will contain singles cells at the applied density of 1 cell/well.
3. All wells were checked one by one after seeding to identify wells containing single cells. In all 13 cases, wells containing single cells were less than the Poisson-prediction. This is reasonable considering that some cells died during the seeding process. Some of these single cells survived and could be further expanded.
4. It is possible that there are wells which were recorded as containing single cells at seeding in fact containing more than 1 cells. Indeed, a small number (up to 8%) of wells which were recorded as not having cells at seeding turned out to have cells later. However, frequency of such overlooking should be small (< 8%). Majority of the recorded single-cell-wells should have contained single cells.
5. No cells from the 3 human tumors survived from the same low-density seeding, suggesting that clonal and low-density growth is a specific feature of dental pulp cells.

Clonal growth or the survival potential at low density is a critical feature of stem/progenitor cells. Therefore, we hypothesized that clonal dental pulp cells will have stronger stem cell features than their parental cells. However, viability and differentiation data of the clonal dental pulp cells in the present study did not support our hypothesis. Some clones had higher viability than the parental cells but there were also some opposite clones. Same apply to differentiation. Consequently, cloning does not seem to provide a strategy for selecting/enriching stem/progenitor cells from human dental pulp cells.

Heterogeneity within a putative stem cell population presents an obstacle for studying their biological features and a challenge to their potential applications. Therefore, effort has been and is still being paid to reduce such heterogeneity. However, diversity may be an intrinsic feature of a cell population where different subpopulations contribute differently. If so, selected and enriched homogeneous cells will become diverse again as they are expanded. Indeed, markers for different differentiation lineages were detected within cells of a colony of bone marrow mesenchymal stem cells [14]. Assuming a single-cell-origin for those colonies, the result would mean intra-clonal heterogeneity.

Intra-clonal heterogeneity would provide an indicator for the limit of homogeneity of a cell population. The differentiation data of clonal cells in our study may suggest some kind of intra-clonal diversity. For example, after induction for adipose-differentiation, some cells had more oil-droplets, some less and others had no. The critical precondition for addressing intra-clonal heterogeneity is the reliability of the single-cell-origin for the clones or colonies being studied. Majority of our clones in the present study should meet this criterion and therefore provide a suitable resource for further comprehensive studying intra-clonal heterogeneity.

## Conclusion

Seeding cells at clonal density into physically separated compartments like wells in a 96 plate and comprehensive observation provides a practical strategy for obtaining clonal cells with increased reliability for their single-cell origin. Dental pulp cell in clones differ from each other and from their parental cells likely indicating that cellular heterogeneity is an intrinsic feature also for human dental pulp cells.

## Acknowledgements

LG: study conception and design, experimental operation, data collection, analysis and interpretation, critical editing of the manuscript. LK: study conception and design, data analysis and interpretation, critical editing of the manuscript. ZZ: study conception and design, experimental operation, data collection and analysis. YM: experimental operation, data collection and analysis. MF and REF: study conception, discussion and critical editing. LG and ZZ were supported by the China Scholarship Council (No.201806370248; No.201806370249). YM was supported by the Hebei eye hospital.

